# *Trithorax* regulates long-term memory in *Drosophila* through epigenetic maintenance of mushroom body metabolic identity and translation capacity

**DOI:** 10.1101/2023.08.08.549887

**Authors:** Nicholas Raun, Spencer G Jones, Olivia Kerr, Crystal Keung, Veyan Ibrahim, MacKayla Williams, Deniz Top, Jamie M Kramer

## Abstract

The role of epigenetics and chromatin in the maintenance of postmitotic neuronal cell identities is not well understood. Here, we show that the histone methyltransferase trithorax (trx) is required in postmitotic memory neurons of the *Drosophila* mushroom body (MB) to enable their capacity for long-term memory (LTM), but not short-term memory (STM). Using MB-specific RNA-, ChIP-, and ATAC-sequencing, we find that trx maintains expression of several non-canonical MB-enriched transcripts, including the orphan nuclear receptor *Hr51*, and the metabolic enzyme *lactate dehydrogenase*. Through these key targets, trx establishes a metabolic state characterized by high lactate levels in MBγ neurons. This metabolic identity supports a high capacity for protein translation, a process that is essential for LTM, but not STM. These data suggest that trx, a classic regulator of cell type specification during development, has an alternative function in maintaining underappreciated aspects of postmitotic neuron identity, such as metabolic state. Our work supports a body of evidence suggesting that a high capacity for energy metabolism is an essential cell identity characteristic for neurons that mediate LTM.

## Introduction

Brain function depends on the coordinated activities of many different cell types. The cell identity of postmitotic neurons is maintained by different combinations of transcription factors (TFs), termed terminal selectors, which activate the expression of effector gene networks^1, 2^. Effector genes define critical neuron identity features such as neurotransmitter type, and synaptic surface receptors. Terminal selectors that maintain postmitotic neuron identity have been elucidated in the simple and well defined nervous system of *C. elegans*^3–6^, and in more complex nervous systems of *Drosophila* and mice^7–12^. While some terminal selectors are required to maintain an array of general cell identity features^13, 14^, others have more specific roles in the activation of specific aspects of neuronal identity^15, 16^. For example, the TF dimmed (dimm) is required specifically to maintain the expression of genes required for secretory function in *Drosophila* secretory neurons^15, 17, 18^. We still lack a complete understanding of the factors that maintain different aspects of neuron identity in a complex nervous system. In addition, the different types of cell identity features that are important for specialized neural functions are not fully elucidated.

Epigenetic modification of chromatin structure has been proposed as a mechanism through which cells establish and maintain their cell identity^1, 19^. The Trithorax group (TrxG) proteins are among the most well-studied epigenetic regulators. TrxG proteins counteract Polycomb group (PcG) proteins to activate the expression of homeotic (hox) genes, which define body segment identity during development^20^. While the role of TrxG and PcG proteins in developmental cell type specification is well defined, their function in maintaining postmitotic cell identity is not extensively explored.

The namesake of the TrxG genes, the *Drosophila melanogaster trithorax (trx)* gene, encodes a histone methyltransferase that methylates histone H3 on lysine 4 (H3K4), a chromatin mark associated with active gene expression. In *Drosophila* there are three enzymes that catalyze H3K4 methylation, trx, trr and Set1^21^. While trr and Set1 regulate bulk levels of H3K4 mono-and tri-methylation, respectively, trx seems to have a more selective role in modification of specific loci^21, 22^. In humans, heterozygous mutations in the orthologous H3K4 methyltransferases (trx=KMT2A/B, trr=KMT2C/D, Set1=SETD1A/B) cause neurodevelopmental disorders (NDDs), such as autism and intellectual disability^23–28^. Individuals with heterozygous mutations in H3K4 methyltransferases do not generally have morphological brain abnormalities^24, 25, 27–30^, suggesting that their cognitive deficits are caused by dysfunction of postmitotic adult neurons. Therefore, in this study we investigated the function of trx in adult postmitotic memory neurons of the *Drosophila* mushroom body (MB). Our data reveals an unexpected role for trx in epigenetic maintenance of MB metabolic state, which is emerging as an essential aspect of postmitotic cell identity for neurons that mediate LTM^31–34^.

## Results

### trx is required in adult MB-neurons for LTM, but not STM.

We investigated the function of the three known *Drosophila* H3K4 methyltransferases Set1, trr, and trx, in the well characterized memory neurons of the *Drosophila* MB^35^. The MB is required for short-and long-term memory (STM and LTM) in flies, but not for learning^36–39^. The MB-specific *R14H06-Gal4* driver^27, 40–42^, was used to drive expression of two unique RNAi lines for each *Drosophila* H3K4 methyltransferase. MB-specific knockdown (KD) flies were then tested for STM and LTM using the classic *Drosophila* memory assay, courtship conditioning^39, 43, 44^. Our previous work showed that Set1 is required in the MB for courtship STM and LTM^27^ and that trr is required for STM (**Fig 1A**)^45^. Here we show that trr is also required in the MB for LTM, while trx is only required for LTM, not STM (**Fig. 1A and S1A**).

**Figure 1.**
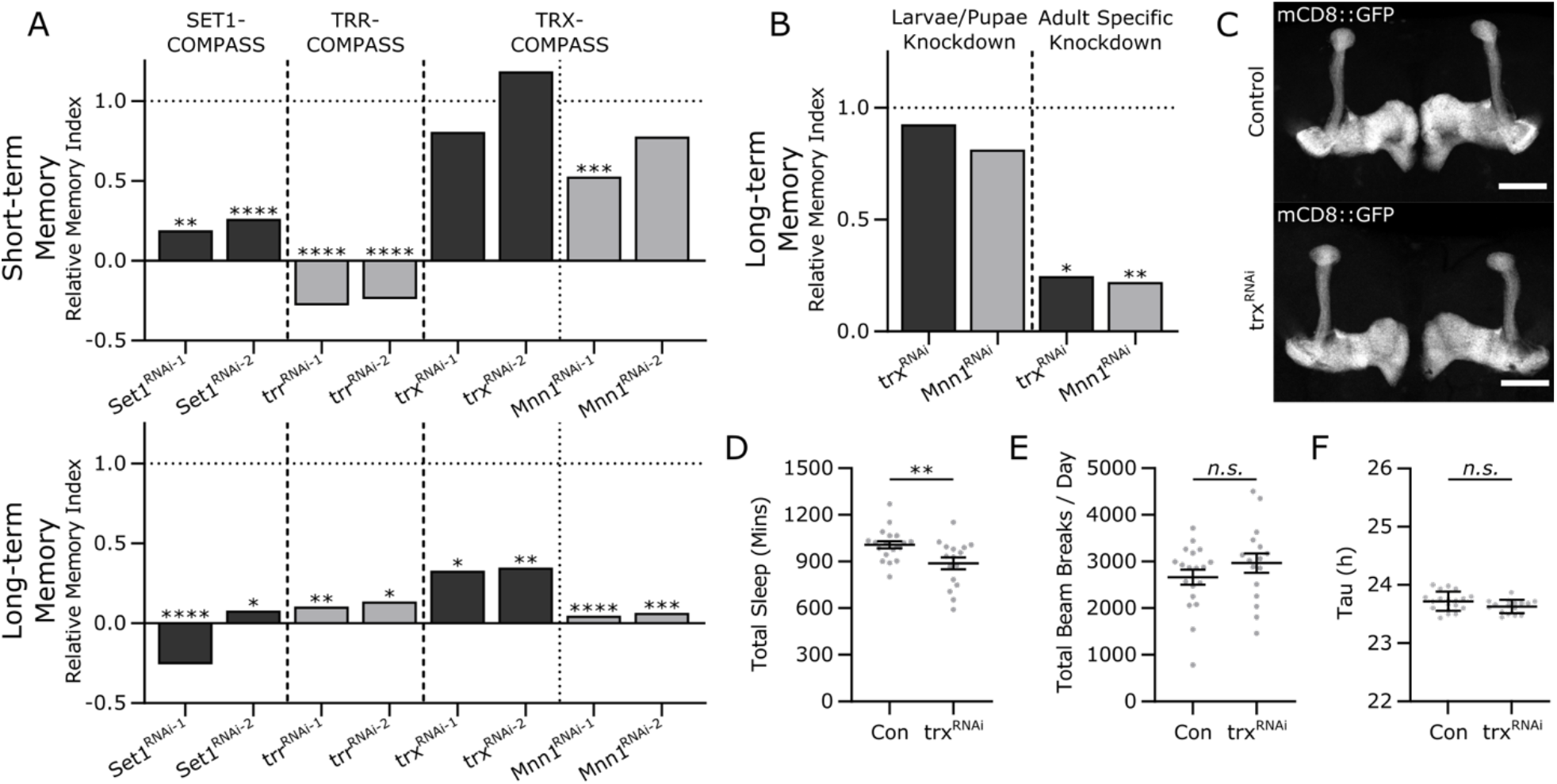
trx is required in the MB for long term courtship memory. **(A)** Bar graphs showing relative memory indices (MI) for courtship STM (upper panel) and LTM (lower panel). Two RNAi lines were used for each gene: *Set1*, *trr*, *trx*, and *Mnn1*. The type of COMPASS complex they are associated with is indicated above. Previously published data for Set1(ref) is shown for comparison. Underlying data can be found in **Fig. S1A**. **(B)** Bar graphs showing LTM upon developmental stage specific MB KD of *trx* and *Mnn1* (Larvae/pupae KD – left panel, adult KD – right panel). Underlying data can be found in **Fig. S1B**. Statistical significance in **(A)** and **(B)** was determined using a randomization test with 10,000 bootstrap replicates. Horizontal dotted line indicates the relative levels of control MIs. **(C)** Confocal Z-stack projections of *trx*-KD MBs compared to controls, labeled with *UAS-mCD8::GFP* driven by *R14H06-Gal4*. Scale bar indicates 50 microns. **(D-F)** Dot plots showing **(D)** sleep, **(E)** activity, and **(F)** circadian rhythm of *trx* MB KD compared to controls. Statistical significance was determined using Student’s t-test. *n.s.* not significant, \**p*<0.05, \*\**p*<0.01, \*\*\**p*<0.001, \*\*\*\**p*<0.0001.

Each Drosophila H3K4 methyltransferase acts in a unique conformation of the complex of proteins associated with SET1 (COMPASS)^21^. The COMPASS complexes consist of 4 common subunits, and additional complex-specific subunits^21^. trx acts in the TRX-COMPASS complex where it interacts with the specific co-factor Menin 1 (Mnn1)^46–48^. MB-specific *Mnn1*-KD also induced LTM-specific defects, with both RNAi lines inducing a strong reduction in LTM, and only one RNAi line having a mild effect on STM (**Fig. 1A and S1A**).

To understand if LTM loss resulted from a developmental or adult-specific role of *trx* and *Mnn1*, we limited the post-mitotic KD in neurons to either the larval/pupal stage, or the adult stage, using temperature sensitive Gal80 (Gal80^ts^)^49^. When *trx-* and *Mnn1*-KD was limited to the larval and pupal stages, we found that memory was unaffected, however, when the KD was limited to the adult, LTM was lost (**Fig. 1B and S1B**). Gross MB morphology also appeared normal following *trx*-KD (**Fig. 1C**), suggesting a limited role for trx in postmitotic MB-morphogenesis. Additionally, we tested MB specific *trx*-KD flies for other behaviours that are linked to the MB, including sleep and activity^50^. *trx*-KD induced mild decrease in total sleep (**Fig. 1D**), with all bouts of sleep differences occurring during the dark period of the day-night cycle (**Fig. S1C**). However, total activity (**Fig. 1E**) and rhythmicity of sleep (**Fig. 1F**) were normal in *trx*-KD flies. Taken together these data demonstrate that trx and its specific cofactor Mnn1 support the capacity of adult postmitotic MB neurons to form LTM.

### trx is dispensable for memory induced gene transcription

A key difference between LTM and STM is that LTM requires *de novo* gene transcription and protein translation, whereas STM does not^51, 52^. Considering the known role of trx as a transcriptional activator we hypothesized that trx might be involved in LTM training induced gene expression in the MB. We therefore analyzed the MB transcriptome in *trx*-KD and control flies, before and after LTM training. In control flies we observed 152 genes that were induced one hour after LTM training compared to naive flies (**Fig. 2A**). These training induced genes were mostly found to be induced in *trx*-KD flies following training (**Fig. 2A**), and generally, were significantly higher expressed in *trx*-KD MBs compared to controls in both naive and trained conditions (**Fig. 2A**). These 152 genes are enriched for GO annotations related to memory formation, such as synapse organization (GO:0050808), learning or memory (GO:0007611), and signal transduction (GO:0007165) (**Fig. 2B**). This suggests that trx does not play a major role in LTM training induced gene transcription.

**Figure 2.**
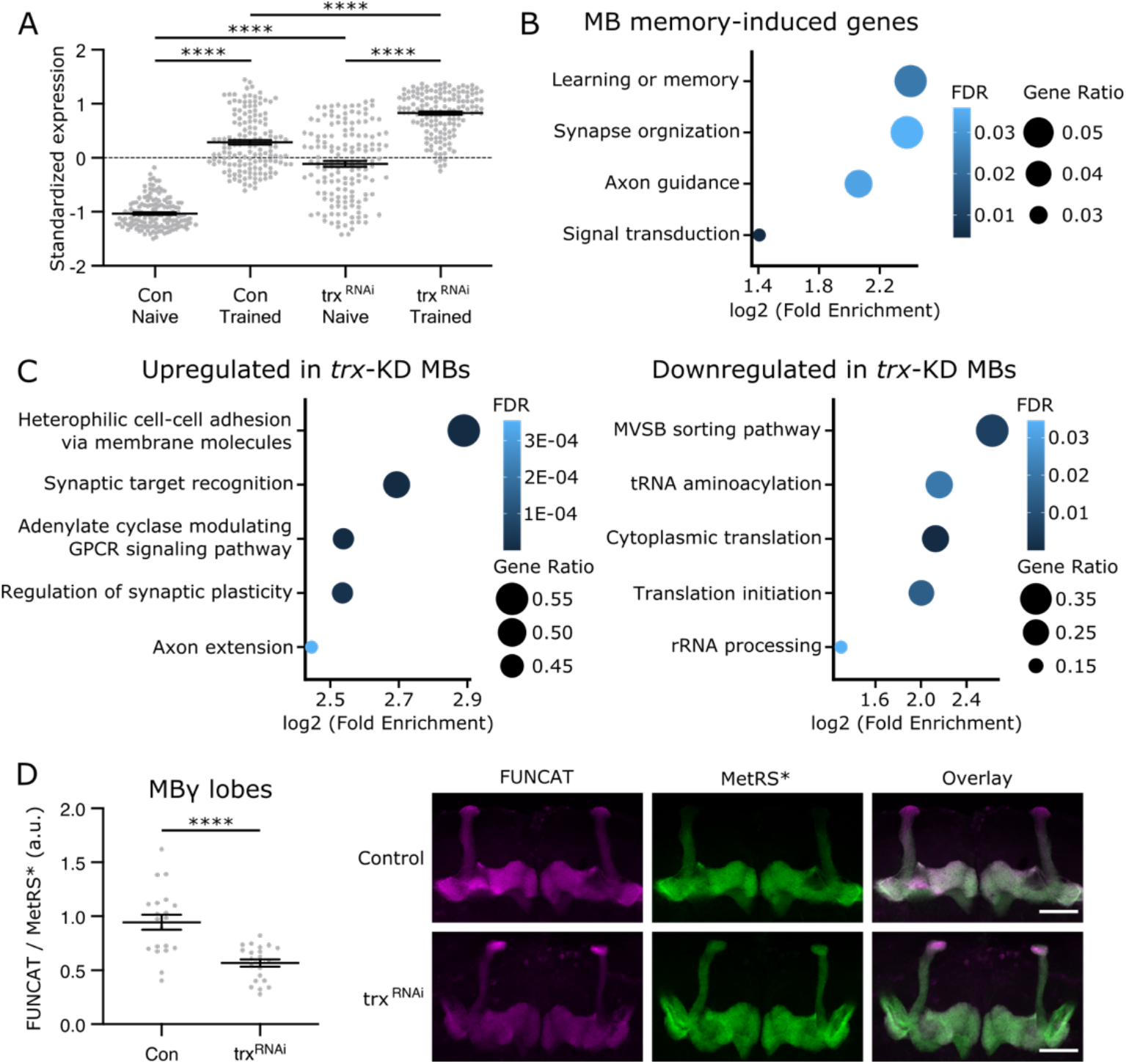
trx regulates translation genes and protein synthesis in MB neurons. **(A)** Dot plot showing standardized expression values for 152 genes that were significantly induced between naïve and trained control flies. These genes are plotted for 4 different conditions: trained and naïve, control or *trx-*KD. The training induced genes identified in controls are expressed at higher levels in *trx*-KD and also show similar induction by training. *P*-values were determined using a one-way ANOVA. **(B)** Bubble plot summarizes gene ontology enrichment analysis of the 152 training induced genes. The 4 displayed terms are the most enriched terms from the 4 representative branches of the GO hierarchy. **(C)** Bubble plot summarizes gene ontology enrichment analysis of up regulated (left panel) and downregulated (right panel) genes when comparing controls to *trx*-KD flies. The 5 most enriched and representative ontologies are shown. For **(B)** and **(C)** dot colour indicates FDR, and size indicates the ratio of genes found in our gene list compared to the reference. Statistical significance was determined using Fisher’s exact test. **(D)** Dot plots showing measurements of FUNCAT fluorescent intensity values normalized to MetRS*::GFP intensity, compared between controls and *trx*-KD in the MBγ lobes. Statistical significance was determined using Student’s t-test. Adjacent representative images show FUNCAT (left panel), MetRS*::GFP (MetRS*) signal (middle panel), and an overlay (right panel). Scale bars indicate 50 microns. \*\*\*\**p<0.0001*

### trx promotes translation in the MB

Since trx does not appear to control LTM training induced gene transcription, we examined other processes that might be compromised in *trx*-KD MBs. Comparing RNA-seq data from control and *trx*-KD MBs (**Table S1**), we found significant upregulation of synaptic genes (**Fig. 2C**), consistent with the observation that LTM training induced genes are overexpressed at the mRNA level in *trx*-KD MBs (**Fig. 2A**). In contrast, downregulated genes were enriched for processes related to translation (**Fig. 2C**). Of the 861 downregulated genes in *trx*-KD MBs, 68 are involved in translation (GO:0006412), including key genes involved in translation initiation (GO:0003743, e.g., eIF2β and 4 eIF2B subunits), 26 structural constituents of the ribosome (GO:0003735) and 11 tRNA synthetase genes (GO:0043039).

We next used fluorescent noncanonical amino acid tagging (FUNCAT)^53, 54^, to quantitatively evaluate translation in the MB. To perform FUNCAT in the MB, a UAS responsive mutant methionyl-tRNA synthetase (*UAS-MetRS\**) was expressed using *R14H06-Gal4*. Expression of MetRS* allows for the incorporation of the non-canonical amino acid azidonorleucine (ANL) into nascent proteins. ANL is not available in the normal fly media and is not incorporated by wild type MetRS, so it is only incorporated into proteins in tissues where MetRS* is expressed after flies are fed ANL-supplemented food. Proteins containing ANL can then be fluorescently labeled using click chemistry^55, 56^ and visualized by confocal microscopy. FUNCAT analysis in the MB revealed that *trx*-KD MBs have significantly lower ANL labeling than controls in the MBγ lobes (**Fig. 2D**), but not the MBα lobes (**Fig. S1D**), providing direct *in vivo* evidence that some regions of *trx*-KD MBs are deficient in translation. Differences in FUNCAT signal were not likely due to differences in ANL dietary intake, since *trx*-KD flies showed no changes in feeding behaviour compared to controls (**Fig. S1E**). Taken together, these results suggest that *trx*-KD MBs have reduced translation capacity in the MBγ lobes, which is likely underlying the observed LTM specific defect (**Fig. 1A**). The reduced translation capacity of trx depleted MBγ neurons may lead to the transcriptional overactivation of LTM training induced genes (**Fig. 2A**) as a compensatory mechanism. Interestingly, the MBγ neurons are the main MB cell type implicated in courtship memory^57–59^.

### Identification of direct trx target genes in the MB

We next sought to understand how trx promotes translation through its function as a H3K4 methyltransferase. We performed ChIP-seq using H3K4me1 and H3K4me3 antibodies on INTACT-isolated MB nuclei from *trx*-KD and control flies. Potential target H3K4me sites in the MB genome were identified by looking for reduction, or loss, of H3K4 methylation peaks following *trx*-KD. We observed 6 significantly reduced H3K4me1 peaks (**Fig. 3A and Table S2**) and 48 reduced H3K4me3 peaks (**Fig. 3B and Table S3**) (FDR < 0.05, Fold change > 1.25). In order to see if trx has the potential to bind at or near these sites in the genome, we used available published trx ChIP-seq data^60^. We overlapped trx binding sites from ChIP-seq with ATAC-seq data that we generated from INTACT isolated MB nuclei to identify regions of open chromatin in the MB genome. This analysis revealed 2288 genes that had trx binding sites within accessible regions of MB chromatin. None of the reduced H3K4me1 peaks were associated with trx-bound genes. Of the 48 genes with reduced H3K4me3 peaks, 22 were associated with trx binding and 10 of these (*MFS3*, *Hr51*, *Dgp-1*, *Ldh*, *JhI-21*, *Ppa*, *Xrp1*, *Gp93*, *Vsx2*, and *CG15747*) also had reduced mRNA levels in *trx-*KD MBs compared to controls (**Fig. 3C and 3D, Table S4**). These data identify a group of genes that are likely activated by trx directly through its role as a H3K4 methyltransferase in the MB.

**Figure 3.**
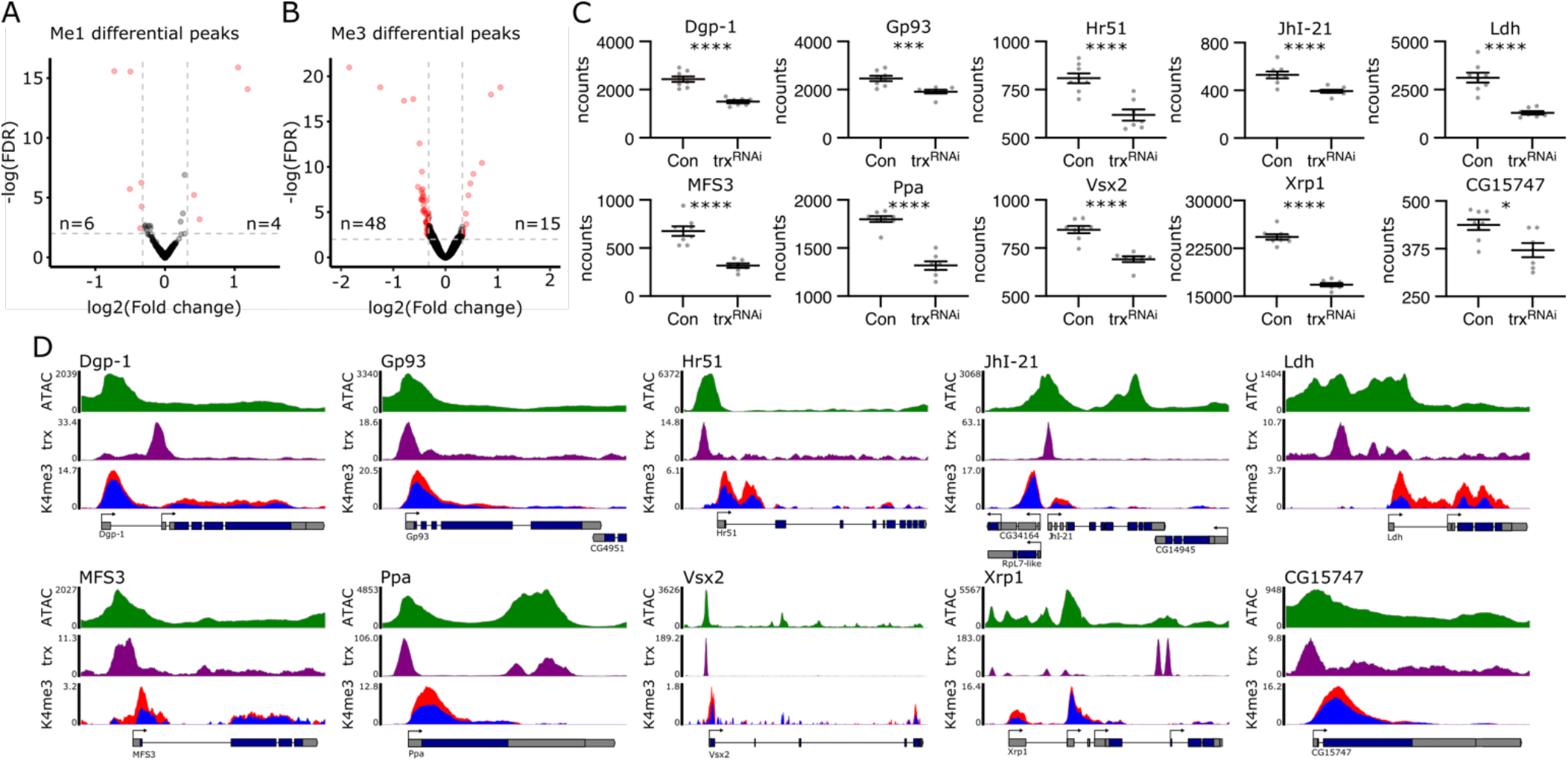
Identification of direct trx target genes in the MB. **(A,B)** Volcano plots showing depleted methylation peaks from differential binding analysis with MB *trx*-KD and control conditions for **(A)** H3K4me1 and **(B)** H3K4me3. Red dots indicate significant peaks with an FDR < 0.01 and Fold change > 1.25, indicated by dashed lines. Statistical significance was determined using a Wald test. **(C)** Dot plots showing the normalized counts (ncounts) between control and *trx*-KD flies for the 10 genes that were found to be significantly downregulated from the 48 reduced H3K4me3 peaks. Statistical significance was determined using a Wald test. **(D)** Genome browser tracks showing chromatin accessibility (ATAC-seq – top panel), trx DNA binding sites (ChIP-seq – middle panel), and H3K4me3 (ChIP-seq – bottom panel) for 10 trx target genes. H3K4me3 tracks display values from both the control (red) and MB-specific *trx^RNAi^* (blue). Below tracks show the different transcripts for each gene in the region compressed with alternative start sites indicated by arrows. \**p*<0.05, \*\*\**p*<0.001, \*\*\*\**p*<0.0001.

### trx direct target genes encompass a set of novel candidate MB identity genes that are required for LTM

Considering the classic known role of trx in cell and tissue specification^61^ we examined whether trx targets may be involved in maintaining the post-mitotic cell identity of MB neurons. To this end, we performed RNA-seq on MB-specific INTACT isolated nuclei and compared it to whole head (WH) nuclear RNA-seq data. Six of the 10 identified trx target genes showed enriched expression in the MB compared to WH (**Table S4**). Four of these, *Dgp-1*, *Hr51*, *Ldh*, and *MFS3*, had greater than 2-fold enrichment of mRNA levels in the MB compared to the WH, similar to what we observe for several canonical pan-Kenyon cell MB identity markers, including; *ey*, *rut*, *Dop1R2*, and *prt* (**Fig. 4A**)^62, 63^. Both the known MB identity genes and the newly identified MB-enriched transcripts (*Dgp-1*, *Hr51*, *Ldh*, and *MFS3*) showed varied levels of expression in the WH but were among the top 50% of expressed genes in the MB (**Fig. 4B**). *Dgp-1* and *Ldh* are among the highest expressed MB enriched genes, similar to the established MB identity genes, *rut* and *prt*. *Ldh* and *MFS3* have among the largest increase in expression from the WH to MB relative to other MB identity genes (**Fig. 4B**).

**Figure 4.**
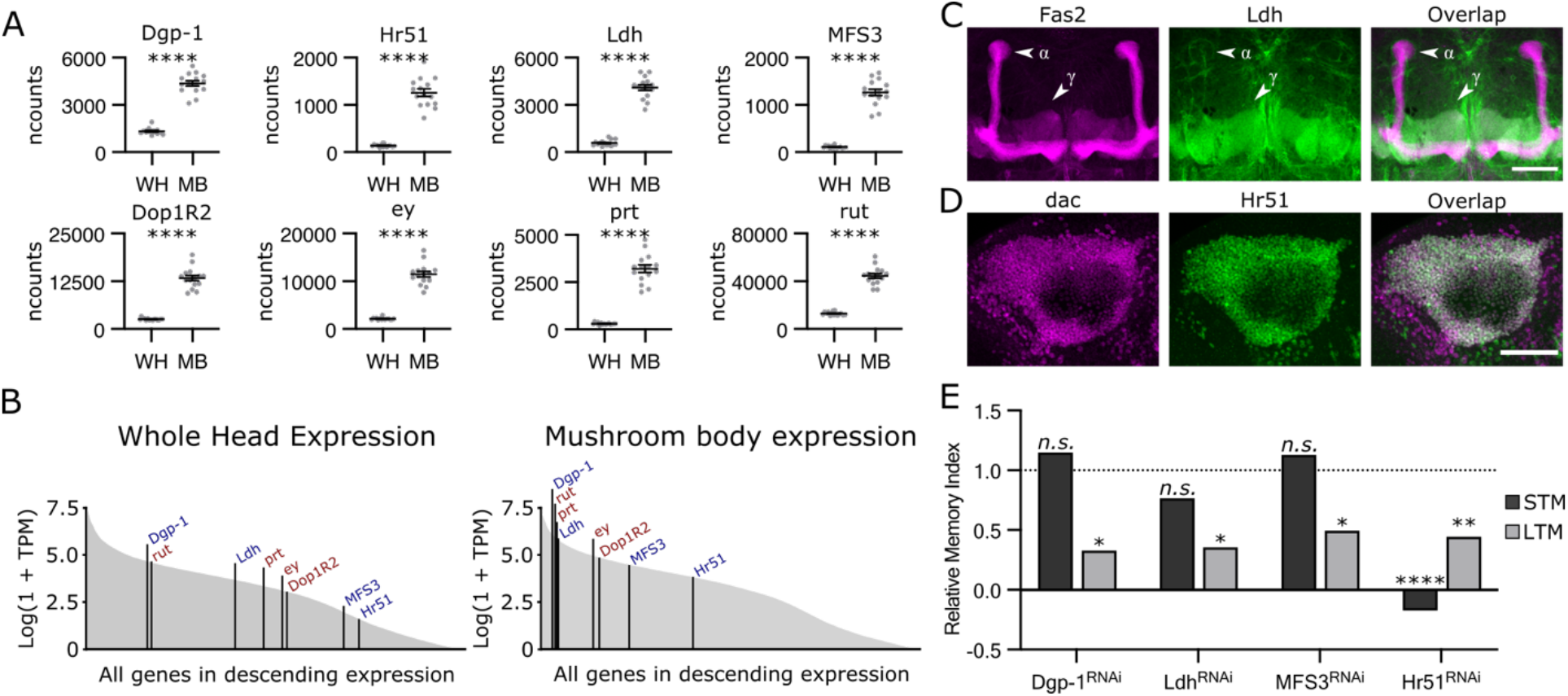
trx target genes are enriched in the MB and required for MB function. **(A)** Dot plots showing the normalized counts (ncounts) of different genes compared between mRNA from the whole head (WH) and from mushroom body (MB) enriched nuclei including trx target genes (upper panel) and 4 canonical MB identity genes (lower panel). Statistical significance was determined using a Wald test. **(B)** Area plots showing the log of transcripts per million (TPM) for all genes expressed in the WH (left panel) or MB (right panel) in descending order. The rank of each trx target gene is marked and labelled in blue, and canonical MB identity genes in red. **(C,D)** Confocal Z-stack projections showing **(C)** MB projections (anti-Fas2 – left panel) and Ldh::GFP (middle panel), or **(D)** MB nuclei (anti-dac – left panel) and Hr51::GFP (middle panel). For **(C)** and **(D)** an overlay is shown (right panel). Scale bars indicate 50 microns. **(E)** Bar graphs showing relative memory indices (MI) for courtship STM (dark grey) and LTM (light grey). Underlying data can be found in **Fig S2E**. Horizontal dotted line indicates the relative levels of control MIs. Statistical significance was determined using a randomization test with 10,000 bootstrap replicates. *n.s.* not significant, \**p*<0.05, \*\**p*<0.01, \*\*\**p*<0.001, \*\*\*\**p*<0.0001

To validate MB enrichment of candidate MB-identity genes we generated or obtained transgenic flies expressing tagged proteins under the control of their native gene regulatory elements for Hr51 (BDSC #38650), Ldh^64^, MFS3^65^, and Dgp-1 (see methods). MFS3 and Dgp-1 were both clearly present in the MB but did not show clear enrichment compared to the surrounding brain tissue (**Fig. S2A and S2B**). In contrast, Hr51 and Ldh proteins both showed striking enrichment in the adult MB. Interestingly, the Ldh protein showed enriched localization to the MBγ neurons and not the MBα/β and α’/β’ neurons (**Fig. 4C**). Ldh enrichment in the MBγ neurons is clear, despite also being expressed in some other neurons and glia throughout the adult brain (**Fig. 4C and S2C**). Notably, the MBγ lobe is the part of the MB that is known to underly courtship memory and was also the region of the MB where translation was compromised in *trx*-KD MBs (**Fig. 2D**)^39, 57^. Ldh has not previously been identified as a MB enriched protein, but was indicated as a candidate MB enriched genes by single cell RNA-seq^62^. Hr51 was clearly localized in dac-positive nuclei of Kenyon cells, with limited expression in the rest of the brain (**Fig. 4D and S2D**). This expression pattern is consistent with previous observations showing nuclear localization in MB neurons^66^. While Hr51 is known to have an important role in MB development and is linked to larval MB neuron identity^67, 68^, its expression and role in the adult MB has not previously been described. These data suggest that some trx target genes may indeed be involved in defining MB cell identity, based on enriched MB expression similar to canonical MB identity genes. Furthermore, the data identifies Ldh as a downstream target of trx that is specific to the MBγ lobe.

To functionally assess the role of these candidate MB cell identity genes we tested their role in courtship memory. Dgp-1, Ldh, and MFS3 are all required in the MB for LTM, but not STM (**Fig. 4E and S2E**). Due to the known role of Hr51 in MBγ neuron remodelling during metamorphosis^68^ we limited *Hr51*-KD to the adult MB using Gal80^ts^ (**Fig. S2F**), The *Hr51* adult specific KD flies showed STM and LTM defects with no effect on MB morphology (**Fig. 4E and S2F**). The role of Hr51 in MB metamorphosis, STM, and LTM suggests that it may have a broader role in the MB compared to trx and its other MB enriched target transcripts, *Dgp-1*, *Ldh*, and *MFS3*, which are only required for LTM.

Taken together, we have identified a group of MB enriched transcripts that are regulated by trx and required in the MB for normal LTM. These non-canonical MB enriched genes are distinct from canonical MB identity genes, and are novel trx target genes, distinct from the known developmental hox target genes.

### trx target genes affect the translation capacity of MB memory neurons

Next, we sought to understand how MB-enriched trx target genes support translation and LTM in the MB. Since Hr51 is a nuclear receptor transcription factor, we reasoned that it may activate important LTM genes downstream of, or in cooperation with, trx. To investigate this, we used a published Hr51 ChIP-seq dataset (ENCSR555TTB)^69, 70^ to identify 4025 genes with Hr51 binding sites that overlap with MB-specific accessible regions, which we generated by ATAC-seq. Among the Hr51 bound genes we identified all four functionally validated MB-enriched trx target genes, *Ldh*, *MFS3*, *Dgp-1*, and *Hr51* itself (**Fig. 5A**). The Hr51 binding sites at these genes directly overlap with MB open chromatin and are adjacent to trx binding sites and differential H3K4me3 peaks (**Fig. 5A and 3D**). This suggests that trx and Hr51 might work together to maintain expression of MB enriched trx target genes, and that Hr51 is self-activating.

**Figure 5.**
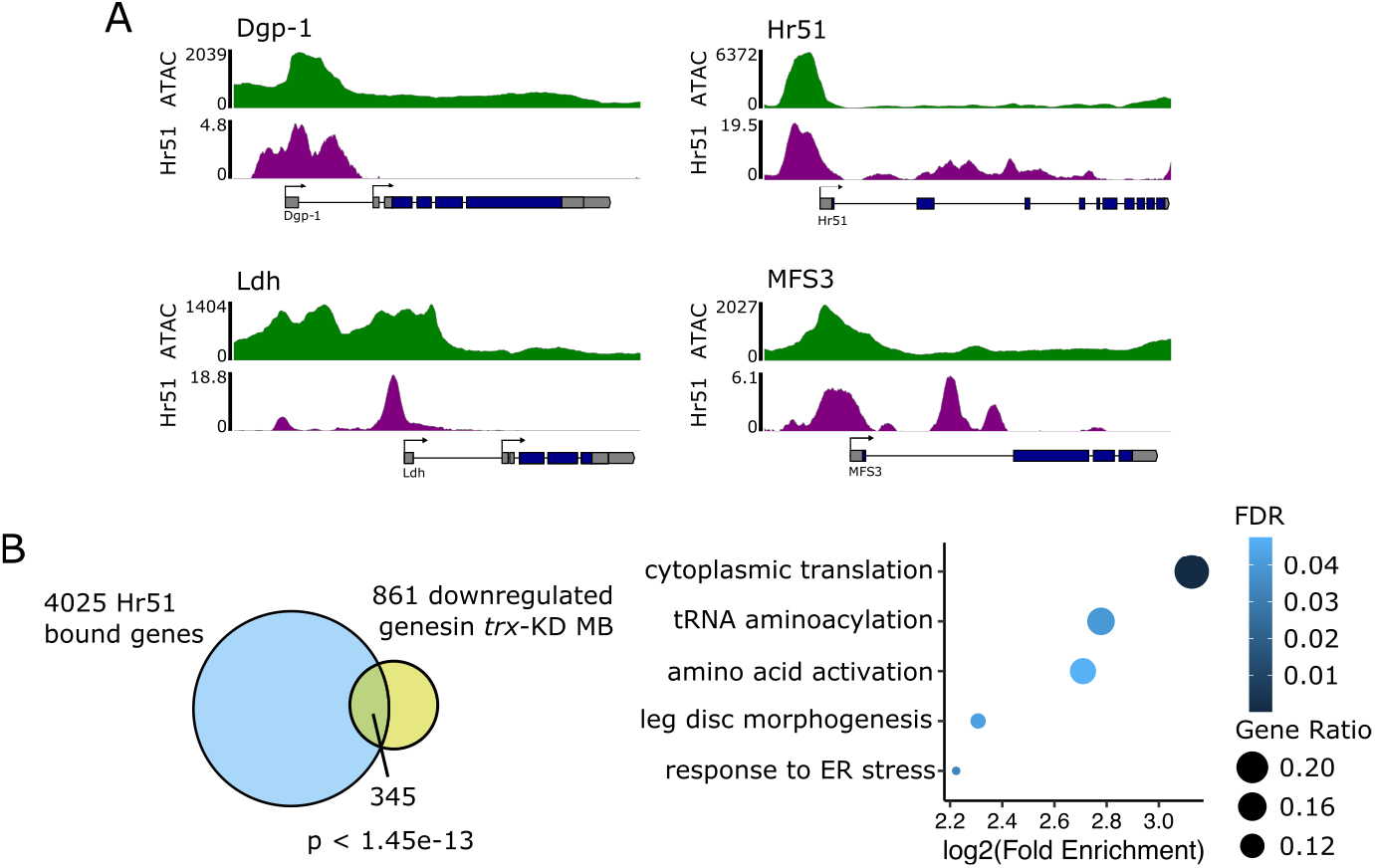
Hr51 target gene analysis. **(A)** Genome browser tracks showing chromatin accessibility (ATAC-seq – top panel), and H51 DNA binding sites (ChIP-seq – bottom panel) for 4 Hr51 target genes. Below tracks show the different transcripts for each gene in the region compressed with major alternative start sites indicated by arrows. **(B)** Venn diagram showing the overlap between Hr51 bound genes and downregulated genes following *trx*-KD. Overlap statistics assume a population of 13,986 coding genes in the *Drosophila* genome. Statistical significance was determined using a hypergeometric test. Adjacent bubble plot summarizes gene ontology enrichment analysis of the overlapping genes. The 5 most enriched and representative ontologies are shown to reduce redundancy. Dot colour indicates FDR, and size indicates the ratio of genes found in our gene list compared to the reference. Statistical significance determined using Fisher’s exact test.

Interestingly, Hr51 bound genes are highly enriched for genes encoding structural ribosomal proteins, including 84 out of 177 genes annotated as structural constituents of the ribosome (GO:0003735). Hr51 bound genes also significantly overlap with genes down regulated in *trx*-KD MBs (p<1.45e-13) (**Fig. 5B**). This includes 395 genes, which are enriched for GO terms related to translation, including cytoplasmic translation (GO:0002181), tRNA aminoacylation (GO:0043039), and amino acid activation (GO:0043038) (**Fig. 5B**). Therefore, it appears that Hr51 may act directly on some trx target genes, including itself, and potentially also acts as a transcriptional activator of the translational machinery independent of trx. Taken together with the observation that Hr51 is required for both STM and LTM in the adult MB, it appears that this orphan nuclear receptor may have a broader role regulating MB identity in a trx-dependent and -independent manner.

We next asked if the other MB enriched trx target genes, *Dgp-1*, *Ldh*, and *MFS3*, might be involved in supporting the translation capacity of MB neurons. Dgp-1 is the ortholog of Gtpbp1, which is required in mouse neurons to resolve ribosomal stalling during translation elongation^71^. In principle, Dgp-1 might influence translation downstream of trx. In contrast, Ldh and MFS3 are not functionally linked to translation. Therefore, we performed FUNCAT on *Ldh*-and *MFS3*-KD MBs to find a possible role in translation. *MFS3-*KD did not affect translation (**Fig. 6A and S2G**). However, translation was compromised in the MBγ neurons upon KD of *Ldh* (**Fig. 6A**), but not in the MBα neurons **(S2G**). Considering that trx and Ldh both are required for MBγ neuron translation capacity (**Fig. 2D and 5D**), LTM, and not STM (**Fig. 1A and 4E**), we tested if Ldh might be a limiting factor in the regulation of LTM downstream of trx. *trx*-KD results in a lower level of Ldh::GFP protein in the MB, similar to that observed following *Ldh*-KD (**Fig. 6B**). To replenish Ldh protein in *trx*-KD MBs, we co-expressed *UAS-Ldh*. Expression of Ldh did restore normal LTM in *trx*-KD MBs, in contrast to control flies expressing *UAS-GFP* (**Fig. 6C and S3A**). These data show that Ldh expression in the MB is a limiting factor for LTM downstream of trx.

**Figure 6.**
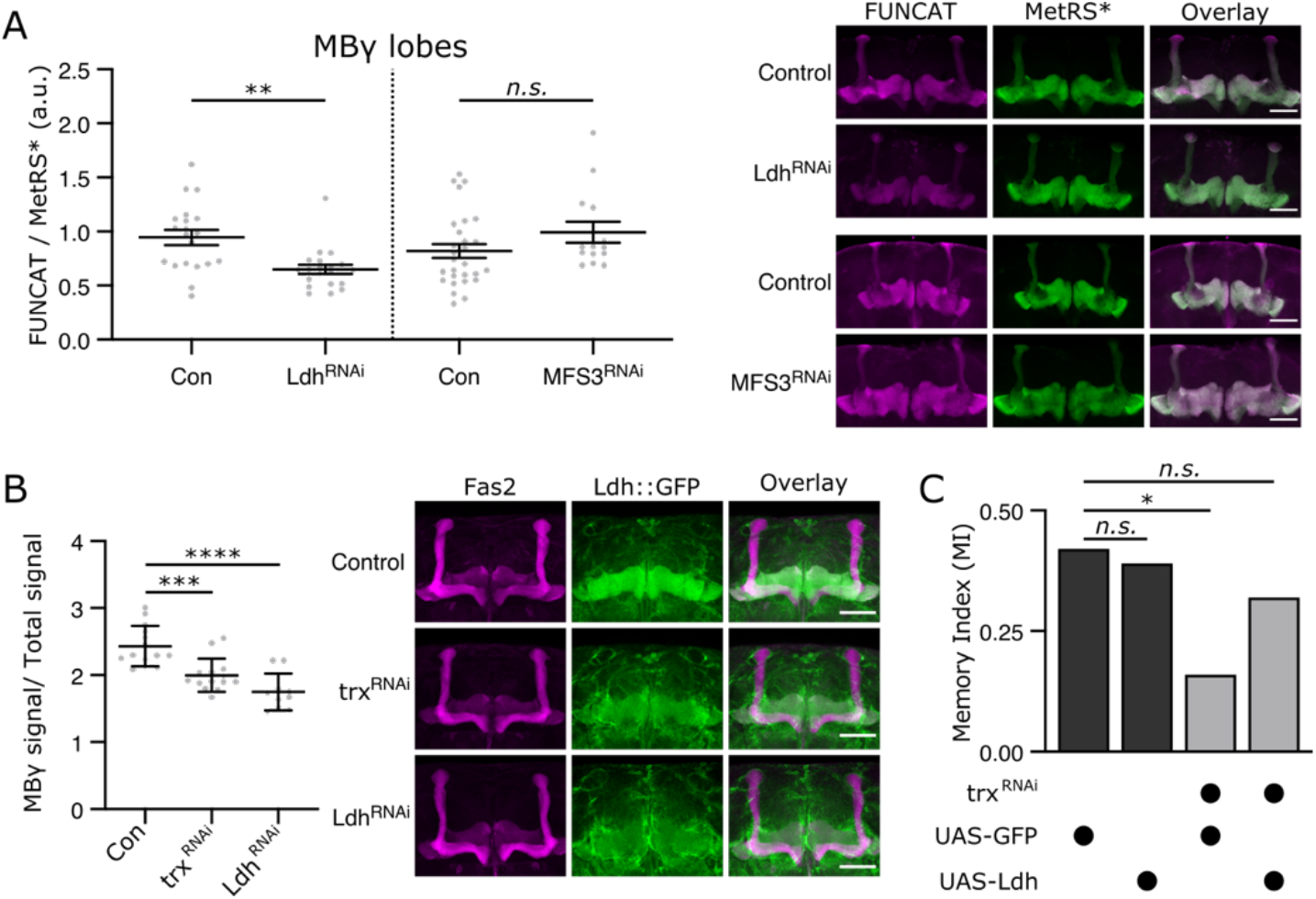
Ldh controls translation downstream of trx. **(A)** Dot plots showing measurements of FUNCAT fluorescent intensity values normalized to MetRS*::GFP intensity, compared between controls and *Ldh* (left panel), or *MFS3* (right panel) in the MBγ lobes. Statistical significance determined using Student’s t-test. Adjacent representative images show FUNCAT (left panel), MetRS*::GFP (MetRS*) signal (middle panel), and an overlay (right panel). Scale bars indicate 50 microns. **(B)** Dot plots show normalized Ldh::GFP signal compared between control, *trx*-KD, and *Ldh*-KD MBγ lobes. Statistical significance determined using Student’s t-test. Adjacent representative images show anti-Fas2 (left panel), Ldh::GFP signal (middle panel), and an overlay (right panel). Scale bars indicate 50 microns. **(C)** Bar graphs showing the memory indices (MI) of different combinations of genetic elements, indicated below, for LTM courtship conditioning. Underlying data can be found in **Fig. S3A**. Statistical significance was determined using a randomization test with 10,000 bootstrap replicates. *n.s.* not significant, \**p*<0.05, \*\**p<*0.01, \*\*\**p*<0.001, \*\*\*\**p*<0.0001

### trx and Ldh maintain a pool of lactate in MBγ lobes

Ldh is required for the reversible conversion of lactate to pyruvate^72^. In mammals, the directionality of this conversion is thought to depend on the isozyme confirmation of LdhA and LdhB tetramers^73, 74^. Unlike in mammals, *Drosophila* only has a single enzyme to perform both reactions, and how the directionality is determined is not well understood^72^. To understand how reduced Ldh protein might impact MB metabolism we assessed levels of lactate and pyruvate in the MB lobes using Gal4 responsive laconic and pyronic FRET sensors, respectively^31, 75^. The laconic and pyronic FRET sensors lose FRET activity upon lactate or pyruvate binding. Brains expressing laconic in the MB that were bathed in 40mM lactate showed a decreased FRET intensity in the MB compared to brains bathed in PBS alone, demonstrating that our methods are able to detect changes in lactate concentration in the MB (**Fig. S3B**). Following both *trx*-and *Ldh*-KDs, we observed that laconic FRET signals increased in the MBγ lobes, indicating that MBγ lactate levels were reduced under these conditions compared to controls (**Fig. 7A**). Pyronic FRET signals decreased in the MBγ lobes following *trx*-and *Ldh*-KD, indicating a corresponding increase in the abundance of pyruvate (**Fig. 7B**). Interestingly, no changes in lactate or pyruvate levels were found in the MBα lobes (**Fig. S3C**), a part of the MB which does not have clear Ldh protein expression (**Fig. 4C**). MFS3 did not impact lactate levels in the MB (**Fig. S3D**). Taken together, these data show that trx and Ldh are required in the MBγ lobe to maintain a pool of lactate.

**Figure 7.**
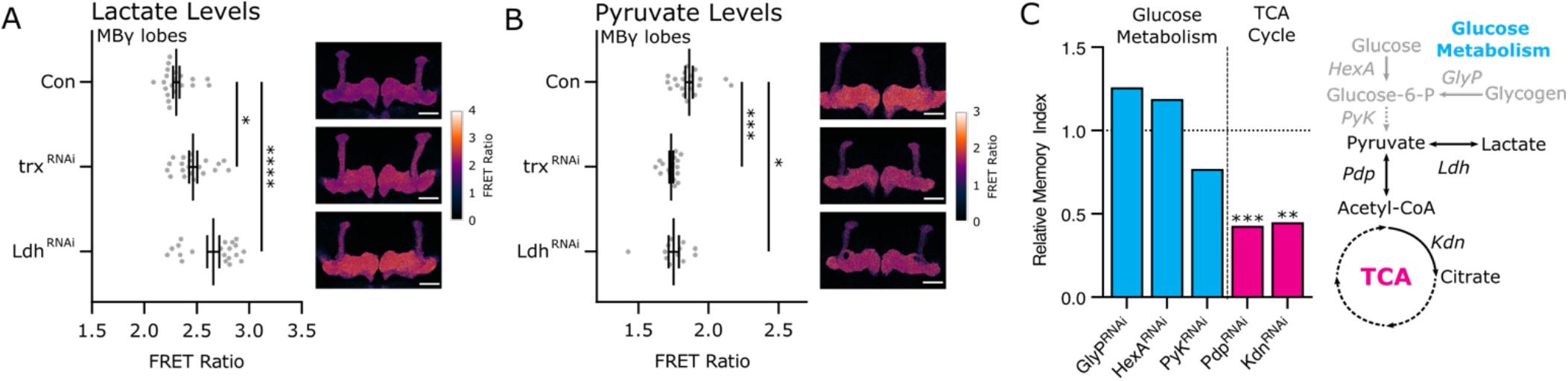
Trx regulates the metabolic state of MBγ lobes necessary for courtship LTM. **(A,B)** Dot plots showing FRET ratio in control, *trx*-KD, and *Ldh*-KD MBγ lobes, measured between CFP and YFP channels obtained after applying a linear unmixing algorithm using **(A)** a laconic FRET sensor or **(B)** a pyronic FRET sensor. Statistical significance was determined using Student’s t-test. Adjacent sample images show differences in FRET ratio corresponding to the control (upper panel), *trx*-KD (middle panel), and *Ldh*-KD (bottom panel) plots. Scale bars indicate 50 microns. **(C)** Bar graphs showing the relative memory indices (MI) of flies with MB specific KD of genes involved in glucose metabolism (blue) and pyruvate’s entry into the TCA cycle (magenta). Horizontal dotted line indicates the relative levels of control MIs. A schematic metabolic pathway highlights the functions of targeted metabolic enzymes involved in glucose metabolism (blue), and entry of pyruvate into the TCA (magenta). Underlying courtship data can be found in **Fig. S3F**. Statistical significance was determined using a randomization test with 10,000 bootstrap replicates.

### Metabolic pathways required for courtship LTM

It has been demonstrated in *Drosophila* olfactory memory that upregulated energy metabolism during memory consolidation is necessary and sufficient for LTM, but not required for STM^31^. LTM has long been thought to be an energy intensive processes, as neuronal activation and LTM formation requires energy intensive processes^76^, including translation and maintenance of membrane potentials^77, 78^. By blocking various critical steps in the metabolism of pyruvate through the TCA cycle, aversive olfactory memory is inhibited^31^. Rationally, courtship conditioning likely also needs this energy to support the taxing translation required for LTM. Nevertheless, the role of metabolism in courtship LTM has not been explored.

To test the role of specific metabolic pathways in courtship LTM, we performed MB-specific KD of critical metabolic enzymes involved in glucose metabolism (GlyP, HexA, PyK), and the entry of pyruvate into the TCA cycle (Pdp and Kdn) (**Fig. 7C**). Glycogen phosphorylase (GlyP) is responsible for the breakdown of glycogen, a carbohydrate storage molecule, into glucose-6-P^79^. GlyP was not required for courtship LTM (**Fig. 7C and S3E**), which was expected, since neurons are not typically thought to store glycogen. Hexokinase A (HexA) is a critical enzyme in glycolysis that facilitates the conversion of glucose into glucose-6-P^80^. Pyruvate kinase (PyK) acts downstream of HexA in the last step of glycolysis to convert phosphoenolpyruvate into pyruvate. Both HexA and PyK were not required for LTM (**Fig. 7 and Fig. S3E**).

For pyruvate to enter the TCA cycle and generate ATP it must be converted into acetyl-CoA by Pyruvate dehydrogenase (Pdh). Pdh is inhibited by Pdh kinase and activated by Pdh phosphatase (Pdp)^81^. Pdp is required for courtship LTM (**Fig. 7C and Fig. S3E**), but not STM (**Fig. S3F**), indicating that entry of pyruvate into the TCA cycle is critical for courtship LTM. Knockdown (Kdn) is a citrate synthase that converts acetyl-CoA into citrate in the first and rate limiting step of the TCA cycle^82^. Kdn was also required for courtship LTM (**Fig. 7C and Fig. S3E**), but not STM (**Fig. S3F**). These results show that glycolysis and glycogen metabolism upstream of pyruvate are not required for courtship LTM, whereas disruption of the components required for pyruvate’s entry into the TCA cycle are especially important for courtship LTM, but not STM. These data support the idea that trx-mediated activation of Ldh might facilitate LTM by enabling high MB energy capacity via the TCA cycle.

## Discussion

trx is a classic epigenetic regulator that targets and activates hox genes to coordinate cell lineage specification during development^61, 83, 84^. However, the role of trx post development in adult tissues, has not been studied. Here, we define a role for trx in the adult MB of *Drosophila*. Loss of trx in the MB causes deficits in LTM and translation capacity, but not STM (**Fig. 1 and 2**). We show that trx is required to maintain mRNA expression of several novel MB-enriched transcripts that have similar expression to known MB cell identity genes (**Fig. 3 and 4**). Based on their expression (**Fig. 4A**), and their role in LTM (**Fig. 4E**), we propose that these are novel MB identity genes that are critical for some, but not all, aspects of MB function. As an example, we show that the trx target gene *Ldh* encodes a protein that is specifically expressed in the MBγ and not the MBα/β neurons. Ldh supports the capacity of MBγ neurons for protein translation (**Fig. 6A**) and is a limiting factor for LTM formation downstream of trx (**Fig. 6C**). trx and Ldh also both contribute to a pool of lactate that is present in the MBγ neurons (**Fig. 7A**). Based on the requirement for TCA cycle enzymes in courtship LTM (**Fig. 7C**), we hypothesize that the MBγ lactate pool might feed the TCA cycle during LTM consolidation. This work identifies an unexpected role for trx in maintenance of metabolic state in MBγ neurons and supports the emerging idea that metabolic state is an important aspect of cell identity in neurons that form LTM.

### Metabolic-state as a trx-dependent aspect of postmitotic neuron cell identity

Neuron identity is defined by a combination of different characteristics, including neurotransmitter type (e.g. cholinergic, GABAergic), morphology, location, and the presence of different types of cell surface receptors^1, 2^. Many neuron types also have specialized functions that require additional specific cellular machinery^2^. Neuron cell identity characteristics are usually established through the expression of a battery of effector genes, often called a subroutine. For example, the cholinergic gene battery includes expression of several enzymes and other specialized proteins to produce acetylcholine (e.g. choline acetyltransferase) and package it into vesicles (e.g. vesicular acetylcholine transporter)^2, 3^. Neuron cell identity features are maintained by terminal selector TFs that continuously activate the expression of the required effector genes^6, 85^. In *Drosophila*, some TFs regulate broad aspects of cell identity^86, 87^, whereas others only activate specific subroutines^15, 17, 18^. One example of this is illustrated in *Drosophila* secretory neurons of the larval brain, where a specific TF, dimm, activates expression of a general neurosecretory subroutine which includes proteins involved in vesicle biology^15, 17, 18^. The full repertoires of cell identity subroutines and the TFs that control them are not known.

Our data suggest that the epigenetic regulator trx may facilitate a metabolic subroutine in MB neurons that supports the capacity of these neurons to form LTM. We show that trx targets several MB enriched transcripts involved in metabolism and translation (**Table S4**). Beyond *Hr51* and *Ldh*, additional MB-enriched trx target genes include *Dgp-1*, potentially involved in translation^71^, metabolite transporters *MFS3* and *JhI-21*^88, 89^, and *Xrp1*, a protein that is involved in DNA damage response and activated in translation deficient cells^90^. Overall, this supports the idea that trx facilitates a MB metabolic subroutine required for LTM formation. Interestingly, it has recently been shown that during LTM consolidation in an aversive olfactory memory assay, the MB, and not the surrounding brain tissues, undergoes a high level of pyruvate flux directed into the TCA pathway for mitochondrial ATP generation^31^. Forced activation of this MB-specific energy influx facilitated the formation of LTM under training conditions that would normally not induce LTM^31^. This suggests that metabolic capacity is a critical feature of MB neuron identity that facilitates the capacity to form LTM. Our findings support this idea by demonstrating that some aspects of MB metabolic state are supported by epigenetic maintenance of MB enriched transcripts.

### Alternate energy strategies genetically encoded in MB subtypes

In *Drosophila* aversive olfactory memory, MBα/β and α’/β’ cells import alanine, which is converted to pyruvate to feed the TCA cycle during LTM consolidation^34^. Here, we show that disrupting the entrance of pyruvate into the TCA cycle also impairs courtship LTM (**Fig. 6D**), suggesting that the requirement for mitochondrial ATP generation may be conserved between olfactory and courtship LTM. However, there appears to be key genetically encoded differences between the MBα/β and α’/β’ neurons that underly olfactory memory, and the MBγ neurons that underly courtship LTM. Specifically, we show here that Ldh protein is expressed in the MBγ neurons but not the MBα/β and α’/β’. Accordingly, *Drosophila* olfactory memory does not appear to require Ldh^34^, while courtship LTM does (**Fig. 4E**). Interestingly, MBγ neurons also have a high lactate to pyruvate ratio compared to MBα/β neurons (**Fig. 7A and B**) and show reduced capacity for translation upon *trx*-KD that is not observed in MBα/β neurons (**Fig. 2D and S1D**).

LTM consolidation in MBα/β neurons requires imported alanine, which is converted to pyruvate as a source of energy^34^. In MBγ neurons, the external energy supply is not yet known. It is possible that MBγ neurons can import alanine as an energy source, however, in contrast to MBα/β neurons, MBγ neurons could also use lactate as an energy source through a one-step conversion to pyruvate by Ldh. The use of lactate as a critical metabolite for *Drosophila* LTM has been proposed, but not clearly demonstrated^91^. In contrast, the use of lactate as a critical neuronal energy source is well studied in mammalian models in a process known as the astrocyte-neuron lactate shuttle (ANLS)^92, 93^. In rodents, astrocytes breakdown stores of glycogen to produce lactate, which is then shuttled to neurons, where it is converted into pyruvate to supply the TCA cycle. Glial export of lactate, and the uptake of lactate in neurons is necessary for rat avoidance LTM^94^, and also supports *de novo* translation in excitatory and inhibitory neurons of the rat dorsal hippocampus^78^. Translation is an energy intensive process which consumes nearly 35% of a cell’s available ATP^95^. It was therefore suggested that the high ATP cost of translation during LTM formation is sustained by increased TCA cycle activity^78^. Our finding that Ldh supports LTM (**Fig. 4E**), translation (**Fig. 6A**), and lactate levels (**Fig. 7A**), suggests a similar mechanism may occur in MBγ neurons.

### Alternative possible uses of lactate for LTM

Here we propose that the lactate pool maintained by trx and Ldh is used to supply the TCA cycle to produce ATP for translation. However, since monitoring lactate flux during courtship LTM consolidation is impossible, we must consider other possible uses of lactate in neurons. It has recently been shown that lactate acts as an agonist for the G-protein coupled HCAR1 receptor, and can modulate neuronal activity in mammals^96, 97^. It is therefore possible that lactate may contribute to LTM through a role in signaling within the MB. An additional alternative use of lactate may be for the post-translational modification of lysine residues. Recently it was shown that neural excitation and social stress correspond to parallel changes in lactylated protein levels^98^. Interestingly, protein lactylation appears to preferentially target Histone H1 in neurons, suggesting that lactylation may be an important epigenetic modification. Protein lactylation also correlated with increased expression of *c-Fos*, an early marker for neuronal activation^99^. However, no study has yet identified direct gene targets of histone lactylation in neurons. Considering these findings, it is possible that lactate contributes to LTM formation through mechanisms other than energy metabolism.

### Hr51 is a candidate terminal selector in the adult MB

The role of trx in Ldh expression is likely facilitative, as *trx*-KD does not completely abolish Ldh expression (**Fig. 3C**). Single cell RNA-sequencing from whole *Drosophila* shows that trx is ubiquitously expressed in all tissues^100^, and therefore cannot be instructive for Ldh activation. A question that arises from this is how a ubiquitously expressed factor like trx can have cell type specific functions. Hr51 is one MB enriched gene that may be an instructive factor for adult MB subtype identity. During pupal developmental MBγ neuron axons are entirely pruned back and then re-extend to form the adult MBγ lobe^67, 68^. Without Hr51, MBγ neuron remodelling can stall, causing the final adult neuron to have an incorrect morphology. Key neuronal identity markers are also lost in adult MB neurons in the absence of Hr51, including trio (specific for MBα’/β’ and MBγ), and Fas2 (specific for MBα/β and MBγ neurons)^68^. In addition, the data presented here shows that Hr51 has other hallmarks of terminal selectors^1^. With Hr51 ChIP-seq data, it was shown that Hr51 has the capacity for autoregulation by binding to its own promoter, and binding to the promoter of additional MB enriched transcripts identified in this study (**Fig. 5A**). Hr51 protein is also highly and very specifically expressed in the nuclei of MB neurons, and in few other cells of the central brain (**Fig. 4D**)^66^. In this study, courtship conditioning revealed that Hr51 is required in adult MB neurons to facilitate STM and LTM (**Fig. 4E**)^68^, suggesting that it likely has a more broad role than trx, and other trx target genes in regulation of MB identity.

## Methods

### Drosophila stocks and genetics

Unless otherwise stated flies were reared on a standard medium (cornmeal-sucrose-yeast-agar) at 25°C and 70% humidity with a 12h:12h light/dark cycle. *Drosophila* stocks were acquired from the Bloomington *Drosophila* Stock Center (BDSC), the Vienna *Drosophila* Resource Center (VDRC), the Kyoto Stock Center (KSC), or donated from other labs. *R14H06-Gal4* flies express Gal4 under the control of a MB specific enhancer for the adenyl cyclase gene *rutabaga* (BDSC #48677) ^42^. UAS-RNAi stock lines used in this study include: *trx^RNAi^* (1=VDRC #37715 & 2=BDSC #31092), *Mnn1^RNAi^* (1=VDRC #17701 & 2=VDRC #110376), *trr^RNAi^* (1=BDSC #29563 & 2=VDRC #110276), *Ldh^RNAi^* (VDRC #31192), *MFS3^RNAi^* (VDRC #330237), *Hr51^RNAi^* (BDSC #39032), *Dgp-1^RNAi^* (VDRC #27490) *GlyP^RNAi^* (VDRC #109596), *HexA^RNAi^* (VDRC #104680), *Kdn^RNAi^* (VDRC #26301), *Pdp^RNAi^* (VDRC #31661), *PyK^RNAi^* (VDRC #49533). *Set1^RNAi^* data was previously published and shown here for comparison purposes only ^27^. *UAS-Unc84::2xGFP* flies were a gift from Lee Henry^101^. Fluorescent labelling of MB cell membranes was done with *UAS-mCD8::GFP* (BDSC #5130 & #5137). *UAS-MetRS*::GFP* flies were a gift from Elaheh Soleimanpour^54^. Flies with *UAS-Laconic* and *UAS-Pyronic* FRET sensors were a gift from Pierre-Yves Plaçais^31, 102^. Fluorescently tagged proteins used in this study include: Ldh::GFP, gifted by Erica Greisbrecht^64^, *MFS3::YFP* (KSC #118654), *Hr51::GFP* (BDSC #38650), and *Dgp-1::GFP* (this paper). The Dgp-1 tagged protein was generated using the CH322-96L23 bacterial artificial chromosome (BAC) containing the *Dgp-1* gene (chr2R from 18165512 to 18183054) was selected from the P[acman] BAC library^103^. This BAC was incorporated into SW102 *E. coli* cells. A multi-tag including SGFP, 3xFLAG, and V5 was amplified from FlyFos022810 DNA. Recombineering was done to insert the multi-tag directly before the stop codon at the end of the *Dgp-1* gene within the CH322-96L23 BAC^104^. This construct was verified through Sanger Sequencing. The modified BACs were then isolated and sent to Genome Prolab for the creation of a transgenic fly with the tagged *Dgp-1::2XTY1::SGFP::V5::preTEV::BLRP::3xFLAGdFRT* transgene.

### Courtship conditioning assay

Short-term memory (STM) and long-term memory (LTM) was assessed using courtship conditioning, as previously described^27, 43, 44^. Briefly, for each fly pair a courtship index (CI) was calculated, which is the proportion of time spent courting over 10 min. The memory index (MI) represents the percentage reduction in courtship behaviour in trained flies compared to naive and is used to compare memory between different genotypes. MI was calculated using the formula: MI = (x̄ CI_naïve_ - x̄ CI_trained_) / x̄ CI_naïve_. Statistics were generated as previously described, using a randomization test with 10,000 bootstrap replicates^27, 44^. For time-course specific experiments, the expression of the RNAi was restricted with Gal80^ts^ (BDSC #7019) by keeping flies at 18°C, or activated by transferring flies to 29°C.

### Immunohistochemistry

Brains were dissected in PBS and fixed with ice cold 4% paraformaldehyde for 30-45 minutes. For immunohistochemistry, fixed brains were blocked in 5% NGS then incubated overnight with the primary antibodies anti-GFP (1:100, Invitrogen: G10362), anti-Fas2 [1:25, Developmental Studies Hybridoma Bank (DSHB): #1D4], or anti-dac (1:100, DSHB: mAbdac2-3), and secondary anti-bodies AlexaFluor 488 or 594 (1:300, Invitrogen: A1108 & A1105). Brains were then mounted in SlowFade Antifade (Invitrogen: S36972) or VectaShield (VectorLabs: H-1900) before imagining. Images were acquired using a Zeiss LSM 510, 710 or 880 confocal microscopes. Confocal stacks were processed and quantified using ImageJ software^105^.

### Sleep, Circadian Behaviour, and activity analysis

The activity assay and sleep analyses were conducted as previously described^27^. Briefly, flies (0-3 days old) entrained in light/dark cycles (LD; 12h:12h) for 3 days are loaded into cuvettes with solid fly food at one end. Flies are monitored for locomotion using Drosophila Activity Monitor 5M (DAM5M) (TriKinetics Inc, MA USA) at 25°C LD for two full days and subsequently in constant darkness (DD) for seven days, totalling nine full days of activity recording at 1-minute resolution. In calculating sleep behaviour in LD, 5 minutes of continuous inaction was considered a sleep bout^106, 107^ and quantified using locally written python code. Circadian behaviour in DD was analyzed using the ActogramJ plug-in in ImageJ^108^ using Lomb-Scargle.

### Gene Ontology Enrichment analysis

Gene Ontology (GO) enrichment analysis was done using PANTHER (version 17.0)^109–112^. For the different sets of gene lists, biological processes were considered with an adjusted p-value cut-off of 0.05, using Fisher’s exact test with FDR multiple test correction. For genes that were induced after training, we used the most enriched term from the hierarchical branches. For all other GO term analyses, only the most enriched terms with between 25-200 genes in the reference genome were included. Semantically redundant terms were also excluded, including only the most significant term.

### Fluorescent noncanonical amino acid tagging (FUNCAT)

Flies were prepared for ANL labeling as previously described^54^. In brief, adult flies were reared on 10mM ANL supplemented sucrose-agar food for 4 days before dissections. Fixed brains were treated with a FUNCAT-reagent overnight, and then stained with 1:500 anti-TAMRA (Invitrogen: MA1-041) and 1:100 anti-GFP (Invitrogen: G10362). Secondary staining and mounting was done as described above. Brains were imagined on a Zeiss LSM 880 microscope and measurements were taken as a mean intensity from the middle slice of the MBγ lobes.

### Feeding Assay and Analysis

The Activity Recording Capillary Feeder (CaFe) assay is previously described^113^. Briefly, the feeding arena is 3D-printed in acrylic. 1% (wt/vol) agar is placed at the bottom of each well to allow *ad libitum* hydration. Liquid food delivered in borosilicate capillaries (VWR: 53432-706) is composed of 2.5% (w/v) Bacto yeast extract (BD: 212750) and 2.5% (w/v) sucrose (Sigma-Aldrich: 57-50-1) and stored at −20°C. Tracking dye (dodecane: mineral oil, 3:1) (Sigma-Aldrich: 297879 & 330779) containing copper reagent (Sigma-Aldrich: 415286) is placed at the top of the liquid food as a tracking marker. Data was analyzed as previously described^113^. Total food consumed over 24 hours was then summed for each fly across all bouts and compared between genotypes.

### Isolation of nuclei tagged in a specific cell-type (INTACT)

Mushroom body nuclei were isolated with INTACT as previously described from a pool of 50-70 male flies per replicate^40^. Briefly, fly heads expressing Unc84::GFP with the MB specific driver, *R14H06-Gal4*^42^, were ground with a pestle in a 1.5 mL tube, and homogenized with buffer containing 0.3% NP40 in a Dounce homogenizer. Nuclear extract was passed through a 40 nm cell strainer. Unc84::GFP labelled nuclei were then immunoprecipitated from homogenate with 1:200 anti-GFP antibody (Invitrogen: G10362) bound to magnetic beads (Invitrogen: 10004D). 1 mL of purified MB nuclei were used for downstream applications.

### RNA-sequencing and data analysis

*trx*-KD RNA-sequencing data was processed as previously described^40^. In brief, raw sequence reads were trimmed to a minimum base quality of 30 using Prinseq (version 0.20.4)^114^. Trimmed reads were aligned using STAR (version 2.5.3a) to the *Drosophila* genome (BDGP release 6)^115, 116^. Reads that aligned to multiple loci, or to one loci with >4 mismatches, and genes that mapped to *Drosophila* rRNA were also removed. Following alignment, samples had on average 25 million useable reads remaining, with no fewer than 15 million reads in any sample. Gene counts were obtained by HTSeq-count (version 0.7.1) using the default union setting. Y-chromosome and mitochondrial genes, and genes with no reads were removed before differential expression analysis using DESeq2 (version 1.30.1). Normalized counts used for comparing gene expression between conditions were generated using DESeq2 normalization (median of ratios).

### ChIP-sequencing and data analysis

ChIP was performed with True ChIP-seq Kit (Diagenode: C01010132) following manufacturers guidelines with the exception that bead bound nuclei were fixed with 36.5% formaldehyde for 1 min. Fixed nuclei were then lysed for 10 minutes with lysis buffer and DNA was sheared with a Covaris M220 for 10 minutes. Immunoprecipitations were done with anti H3K4me1 and H3K4me3 antibodies (Diagenode: C15410194 & C15410003). DNA was purified using MicroChIP DiaPure columns (Diagenode: C03040001), and libraries were generated with the MicroPlex Library Preparation Kit (Diagenode: C05010001). Completed libraries were sent to Genome Quebec for 100bp paired end sequencing on an Illumina NovaSeq 6000 S4.

Raw sequence reads were trimmed using trimmomatic to remove low quality bases (version 0.39)^117^. Trimmed reads were aligned with Bowtie (version 2.4.1) with the very-sensitive setting to the *D. melanogaster* genome (BDGP release 6)^115, 116^. Reads were then filtered for remove low mapping quality (<4). Reads aligning to mitochondrial chromosomes and scaffolds, were removed. Following alignment, H3K4me3 samples had on average 60 million useable reads remaining, with no fewer than 45 million reads in any sample. H3K4me1 samples had on average 57 million useable reads remaining, with no sample having fewer than 15 million useable reads remaining.

ChIP peaks were next identified with the macs2 (version 2.2.7.1)^118^ callpeak function with a cut-off of *q*<0.01. Peaks were filtered against the ENCODE blacklist to remove peaks occurring in regions with high occurrences of artifacts^119^, and then used to generate a consensus peak set. Differential peak analysis was performed with DiffBind (version 3.0.15)^120^. Peaks were annotated with ChIPSeeker (version 1.26.2)^121^. ChIP data for trx (GSE24521) was obtained from the GEO database^60^. ChIP data for Hr51 (ENCSR555TTB) was download from the ENCODE portal^69^. ChIP and ATAC tracks were visualized using pyGenomeTracks (version 3.6)^122^.

### Metabolite analysis using FRET sensors

Adult male brains expressing FRET sensors in MB neurons, specified by the *R14H06-Gal4* driver^42^, were imaged with a Zeiss LSM 880 to capture three relevant channels simultaneously after excitation at 458 nm. Emission windows of the channels are donor (CFP) channel, 480 – 500 ± 5 nm; acceptor (YFP) channel, 525 – 545 ± 5 nm; and autofluorescence (AF) channel, 595 – 615 ± 5 nm. Quantification was done in the MB as a ratio of the CFP and YFP channels after a previously described linear unmixing algorithm^75^, to correct for background and autofluorescence.

## Data Availability

High-throughput sequencing data is publicly available online at GEO and ENCODE.

## Acknowledgements

We thank Elahah Soleimanpour and Daniela Dieterich for their expert guidance with FUNCAT and Lucía Durrieu and Pablo Wappner for their expert guidance in employing linear unmixing. Thanks to the Bloomington Drosophila Stock and the Vienna Drosophila Resource Center for providing *Drosophila* stocks. This project was funded by a Canadian Institutes of Health Research Project Grant to JMK and a Natural Sciences and Engineering Research Council of Canada Postgraduate Scholarship to NR.

## Notes

### Competing Interest Statement

The authors have declared no competing interest.

