## Supplementary Figures for "*Trithorax* regulates long-term memory in *Drosophila* through epigenetic maintenance of mushroom body metabolic identity and translation capacity"

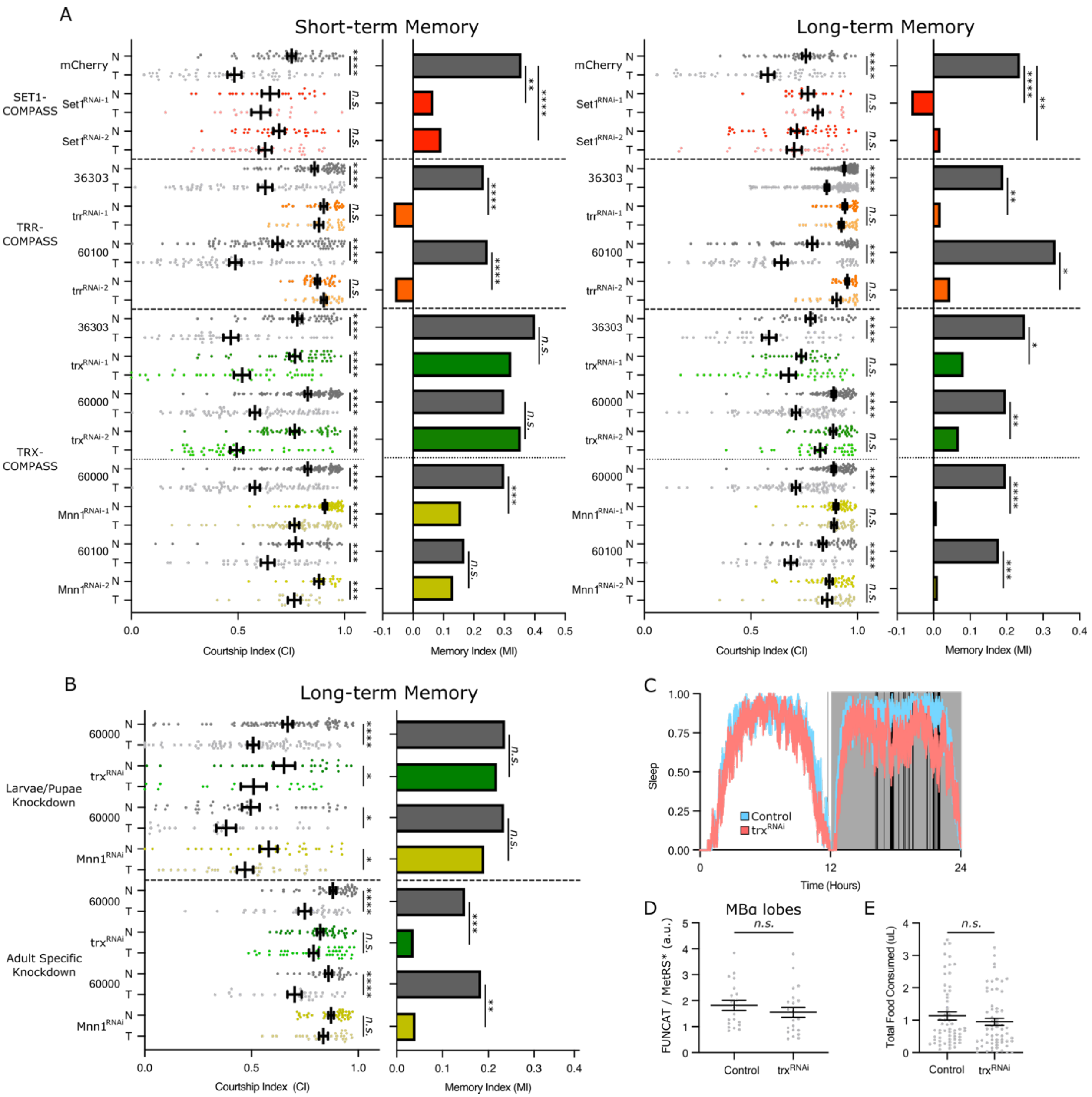


**Supplementary Figure 1**

**(A,B)** Dot plots showing the courtship indices (CIs) for naïve (N) or trained (T) MB KD flies of the COMPASS complex genes. Controls in grey and RNAi in assorted paired colours. Darker coloured dots are naïve and lighter dots are trained. Statistical significance was determined using a Mann-Whitney Test. Adjacent bar graphs show the corresponding memory index (MI) derived from the CI (see methods). Statistical significance between MIs was determined using a randomization test with 10,000 bootstrap replicates. **(A)** Two RNAi lines were used for each gene *Set1*, *trr*, *trx*, and *Mnn1*, indicated left, based on the COMPASS branch they are associated with (grouped by dashed lines). The two components of the trx complex are divided by a dotted line. Courtship STM (left panel) and LTM (right panel) was assessed. **(B)** KDs with one RNAi for trx and Mnn1 were assessed for LTM while KD was limited to the larvae/pupae or to adult flies. **(C)** Ribbon plot showing average sleep per minute of *trx*-KD flies (red) compared to control flies (blue), averaged over 48 hours. The proportion of flies asleep were plotted as 0 (awake) or 1 (all asleep) in 1-minute intervals. Line thickness indicates +/- SEM. White background indicates objective day and dark grey background indicates objective night. Vertical black lines indicate at least 2 contiguous blocks of statistically significant differences in sleep behaviour. Statistical significance was determined using Student's t-test. **(D)** Dot plots showing measurements of FUNCAT fluorescent intensity values normalized to MetRS*::GFP (MetRS) intensity, compared between controls and *trx*-KD in MBα lobes. Statistical significance was determined using Student's t-test. Panels with sample images in **Fig. 2D**. **(E)** Dot plot showing food consumed by control and *trx*-KD flies over a 24-hour period. Statistical significance was determined using Student's t-test. *n.s.* not significant, *$p<0.05$, **$p<0.01$, ***$p<0.001$, ****$p<0.0001$.

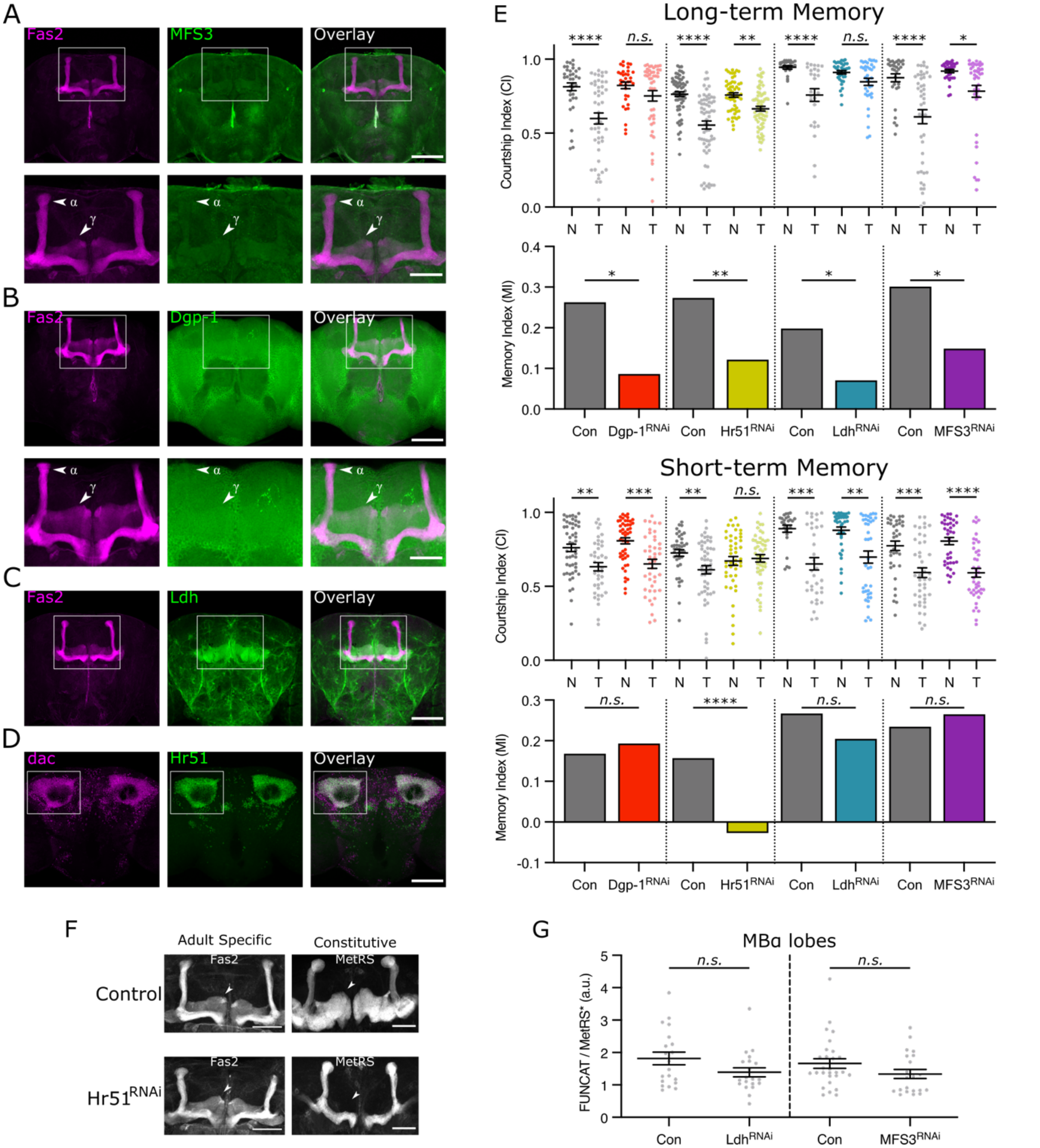


**Supplementary Figure 2**

**(A-D)** Confocal Z-stack projections showing **(A-C)** MB projections (anti-Fas2) or **D** MB nuclei (anti-dac – left panel), **(A)** MFS3::YFP, **(B)** Dgp-1::GFP, **(C)** Ldh::GFP, and **(D)** Hr51::GFP (middle panel), and an overlay (right panel). In **(A)** and **(B)** regions of interest are defined by a white box and shown immediately below. In **(C)** and **(D)** regions of interesting are defined in **Fig. 4C** and **4D**. Scale bars indicate 50 microns. **(E)** Dot plots show the courtship indices (CIs) for naïve (N) and trained (T) MB KD flies of trx-target genes that were assessed for LTM (upper panel) and STM (lower panel). Controls in grey and RNAi in assorted paired colours. Different RNAi are grouped with controls and divided by a dotted line. Darker coloured dots are naïve and lighter dots are trained. Statistical significance was determined using a Mann-Whitney Test. Bar graphs below show the corresponding memory index (MI) derived from the CIs (see methods). Statistical significance was determined using a randomization test with 10,000 bootstrap replicates. **(F)** Confocal Z-stack projections showing control or *Hr51*-KD MBs with adult-specific or constitutive-KD. Brains labelled with anti-Fas2 or MetRS*::GFP (MetRS) to highlight MB projections. Arrow indicates where MBγ lobes should be. Scale bars indicate 50 microns. **(G)** Dot plots showing measurements of FUNCAT fluorescent intensity values normalized to MetRS*::GFP intensity, compared between controls and Ldh (left panel), or MFS3 (right panel) in the MBα lobes. Statistical significance determined using Student's t-test. Sample images are in **Fig. 6A**.

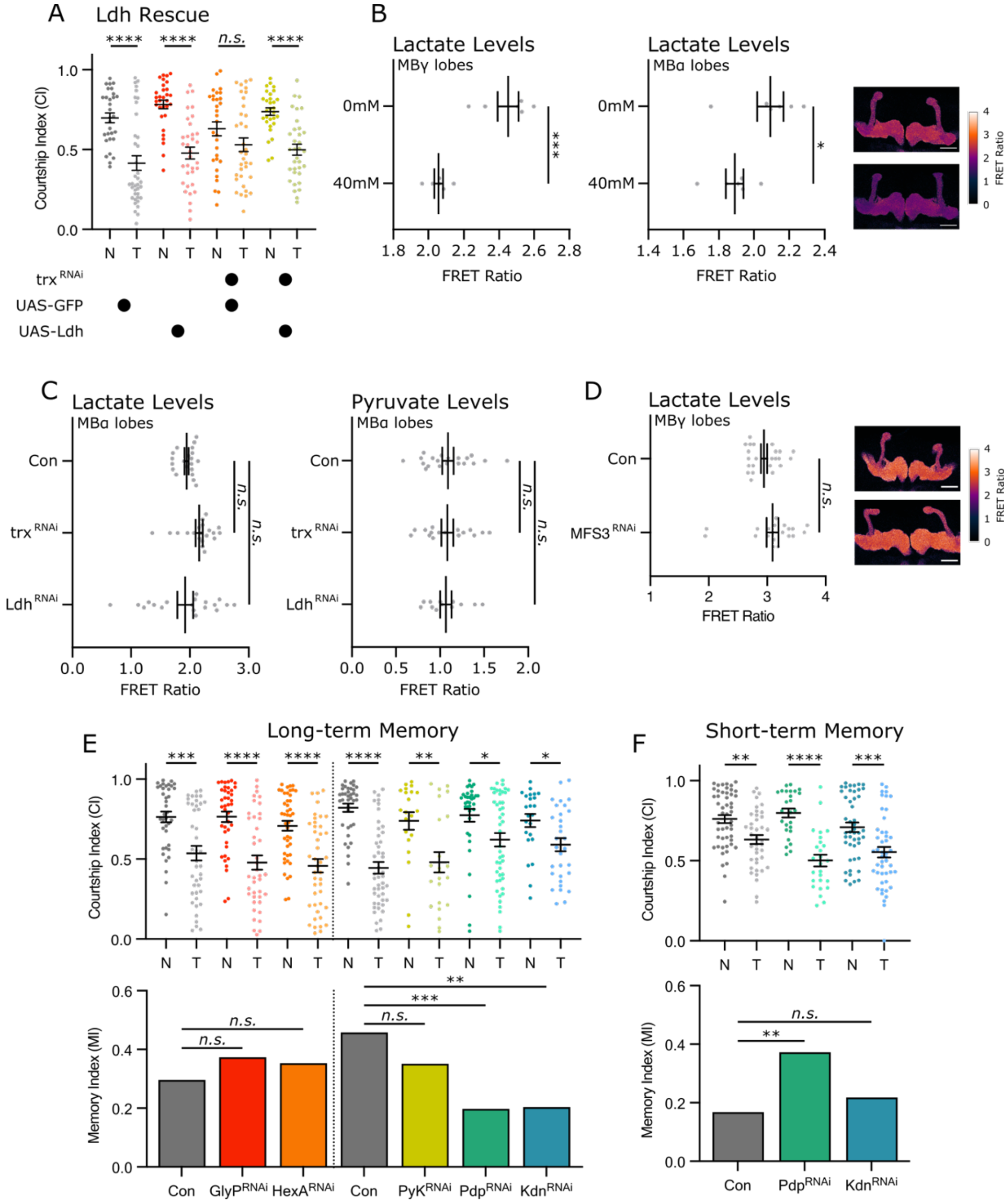


**Supplementary Figure 3**

**(A)** Dot plots showing the courtship indices (CIs) for naïve (N) or trained (T) flies with different combinations of genetic elements (indicated below) expressed in the MB and assessed for LTM. Controls in grey and experimental transgenes in assorted paired colours. Darker coloured dots are naïve and lighter dots are trained. Statistical significance determined using the Mann-Whitney test. **(B)** Dot plots showing FRET ratio in from MBγ (left panel) or MBα (right panel) lobes following a 20-minute treatment of 0mM or 40mM L-lactate dissolved in PBS. Statistical significance determined using Student's t-test. Adjacent sample images show differences in FRET ratio corresponding to the 0 mM lactate (upper panel) and 40 mM lactate (bottom panel). Scale bars indicate 50 microns. **(C,D)** Dot plots showing FRET ratio using **(C)** laconic (left panel) or pyronic (right panel) FRET sensors with control, *trx*-KD, and *Ldh*-KD in MBα lobes or **(D)** laconic FRET sensors with control and *MFS3*-KD in MBγ lobes. For **(C)** Sample images are in **Fig. 7A** and **B.** For **(D)** adjacent sample images show differences in FRET ratio corresponding to the control (upper panel), and *MFS3*-KD (lower panel). Scale bars indicate 50 microns. **(E,F)** Dot plots show the courtship indices (CIs) for naïve (N) or trained (T) flies of metabolic genes that were assessed for **(E)** LTM and **(F)** STM. Controls in grey and RNAi in assorted paired colours. Different RNAi are grouped with controls and divided by a dotted line. Darker coloured dots are naïve and lighter dots are trained. Statistical significance determined using a Mann-Whitney Test. Bar graphs below show the corresponding memory index (MI) derived from the CI (see methods). Statistical significance determined using a randomization test with 10,000 bootstrap replicates. *n.s.* not significant, $^{*}p<0.05$, $^{**}p<0.01$, $^{***}p<0.001$
